# Human sand fly challenge elicits saliva-specific innate and type 1-polarized immunity that promotes *Leishmania* killing

**DOI:** 10.1101/2025.02.25.640210

**Authors:** Maha Abdeladhim, Clarissa Teixeira, Roseanne Ressner, Kelly Hummer, Ranadhir Dey, Regis Gomes, Waldionê de Castro, Fernanda Fortes de Araujo, George W. Turiansky, Eva Iniguez, Claudio Meneses, Fabiano Oliveira, Naomi Aronson, Joshua R. Lacsina, Jesus G. Valenzuela, Shaden Kamhawi

## Abstract

In *Leishmania*-endemic areas, humans are constantly exposed to sand fly bites. To explore the immune consequences of this chronic vector exposure, we performed a human challenge study with the sand fly *Lutzomyia longipalpis*. Peripheral blood mononuclear cells were collected from fifteen healthy volunteers who underwent multiple controlled exposures to sand fly bites. We identified two *Lu. longipalpis* salivary proteins, LJM19 and LJL143, which elicited type 1 cytokine responses in cells from exposed individuals and which correlated with enhanced killing of *Leishmania* parasites in co-cultured macrophages. Interestingly, LJM19 also exerted this parasite-killing effect in cells from unexposed individuals, consistent with innate immune activation. In support of this, both LJM19 and LJL143 stimulated the production of the innate cytokines IL-1β and IFN-α. Our results demonstrate that repeated exposure to sand fly bites induces innate and adaptive cytokine responses to vector salivary proteins that can be co-opted to protect humans against *Leishmania* infection.

## INTRODUCTION

Leishmaniasis is a neglected vector borne disease caused by protozoan parasites of the genus *Leishmania*, which are transmitted via the bite of phlebotomine sand flies. Each year, over 700,000 new cases of leishmaniasis are reported worldwide.^1^ Sand fly saliva plays a critical role in *Leishmania* transmission. When female sand flies bite their host to take a blood meal, they inject a complex mixture of immunogenic and biologically active salivary proteins into the skin. Collectively, these foreign salivary antigens trigger a host immune response that favors parasite establishment.^2^

In *Leishmania*-endemic areas, humans are constantly exposed to these salivary antigens, resulting in an adaptive immune response which induces durable cytokine responses from peripheral blood mononuclear cells (PBMCs). These cytokine responses can be recalled up to ten years after the most recent sand fly exposure.^3,4^ Similarly in the skin, the ability of sand fly bites to elicit a delayed-type hypersensitivity (DTH) response can persist for decades in humans with ongoing sand fly exposure.^5^ For patients with cutaneous leishmaniasis (CL), their cellular and antibody responses to certain sand fly salivary proteins strongly correlate with CL lesion size and whether the disease is localized or disseminated.^6,7^ This suggests that human immunity to sand fly saliva modulates disease severity and susceptibility to leishmaniasis. Furthermore, while there have been numerous clinical studies on how factors like chronic malnutrition^8^ and helminth infection^9^ affect the immune system, there is only a handful of studies that have investigated how repeated vector bites and exposure to salivary antigens – many of which are immunomodulatory – impact immune responses in humans.^4,5,10,11^

Human immunity to sand fly saliva has been investigated through human vector challenge studies, where healthy volunteers are exposed to the bites of uninfected sand flies. Although only a few such human challenge studies have been conducted, they demonstrate that most volunteers develop a skin DTH response to sand fly bites and a mixed cellular type 1/type 2 cytokine response when their PBMCs are stimulated with salivary gland extract (SGE) *ex vivo*.^4,5,10^ One of the recent human challenge studies extended these findings by identifying individual salivary proteins from the Afro-Asian sand fly *Phlebotomus duboscqi* that induce strong IFN-γ or IL-10 responses in PBMCs.^10^

In this human sand fly challenge study, we investigated cellular immunity to saliva of the sand fly *Lutzomyia longipalpis* (*Lu. longipalpis*), the principal vector of visceral leishmaniasis in the Americas, in individuals exposed to uninfected *Lu. longipalpis* bites multiple times over several months. Our objective was to simulate the repeated exposure to sand fly saliva that occurs in *Leishmania*-endemic populations. We identified two sand fly salivary proteins that elicit robust innate and type 1-polarized cytokine responses in human immune cells and enhance killing of *Leishmania* parasites *ex vivo*.

## RESULTS

### Characteristics of study participants

Table 1 presents the demographics, allergy history, and baseline plasma IgE levels of the study participants. Volunteers were screened for allergy status and plasma IgE to exclude individuals potentially at higher risk for developing severe allergic reactions to sand fly bites. Fifteen healthy volunteers were enrolled in the study. The median age was 27 years (range 22 to 46), and the majority were male (12 of 15, 80%). Figure 1 provides a schematic of the study design. Loss to follow-up began at exposure #6, primarily due to active duty military participants being transferred out of the study area (*n* = 4) and one participant being discharged from the military. Three participants halted their participation due to study-related reactions to sand fly bites, which included large areas of urticaria, pruritus, and erythema, raising concerns about potentially worsening symptoms with future sand fly exposures. The number of fed sand flies was high and consistent across all feedings, with an average of 8 to 9 fully fed and 0 to 1 partially fed sand flies out of 10 total per participant at each exposure.

**Fig 1.**
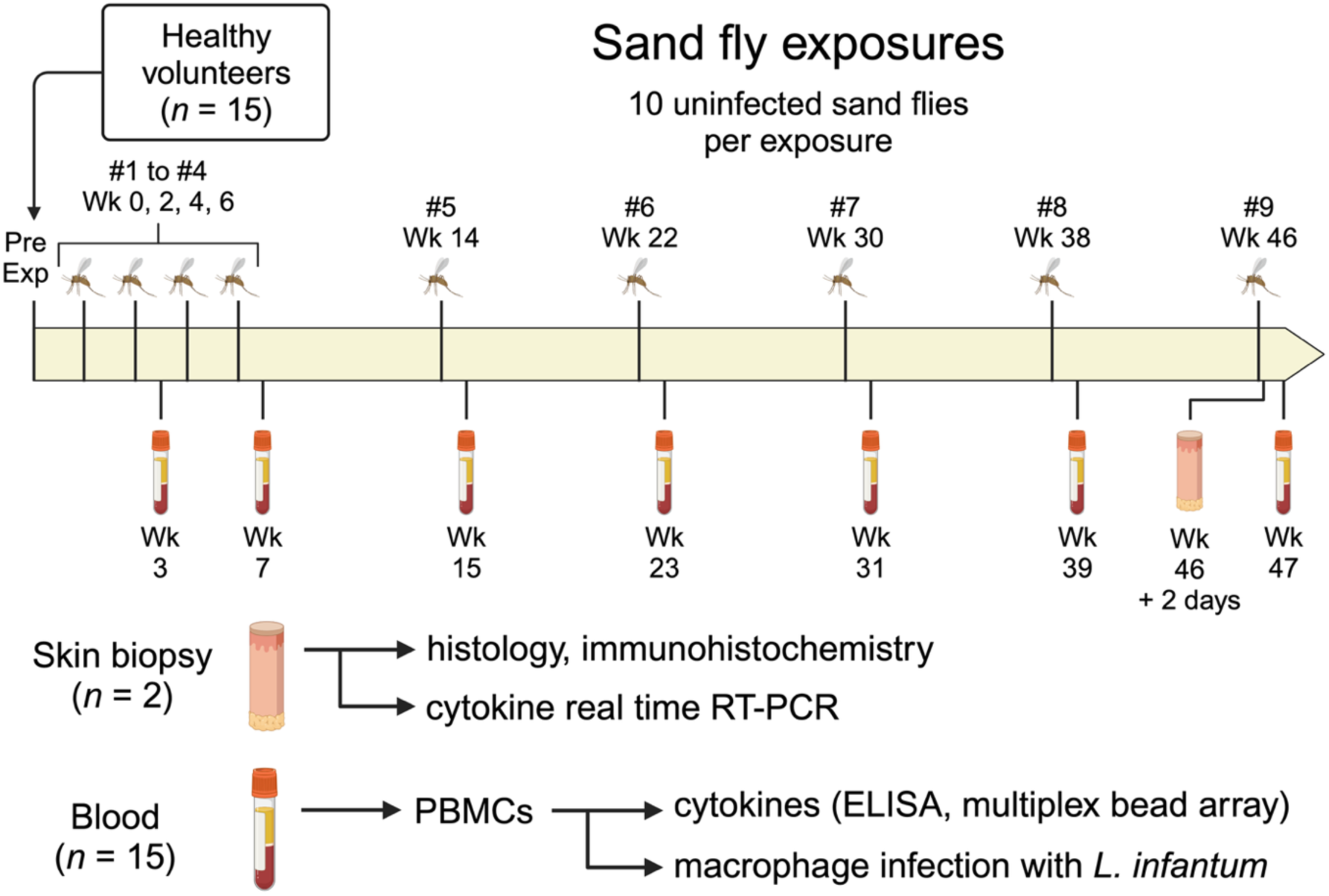
Schematic of clinical study. Healthy volunteers (*n* = 15) were exposed to the bites of uninfected *Lutzomyia longipalpis* sand flies once every two weeks for exposures #1 through #4, then once every eight weeks for exposures #5 to #9. Blood was collected ∼1 week after exposures #2, #4, and #5 to #9, followed by PBMC isolation to perform the indicated assays. Skin punch biopsies were obtained from two participants 48 hours after exposure #9, with one biopsy from a sand fly bite site and one biopsy from the contralateral arm for each participant. *Pre Exp*, pre-exposure study visit. Created in BioRender.com.

**Table 1.** Study participant characteristics.

| Participant ID | Completed Sand Fly Exposure # | Age | Sex | Race | Allergy History | Baseline Plasma IgE (kU/L) | Reason for Early Withdrawal or Skipped Feedings |
| --- | --- | --- | --- | --- | --- | --- | --- |
| 1 | 9 | 26 | M | White | None | 17.9 | - |
| 2 | 5 | 26 | M | White | None | 9.9 | Military transfer orders |
| 3 | 6 | 26 | M | Asian | Seasonal allergies<br>Asthma | 114.0 | Discharged from military |
| 4 | 8 | 40 | M | White | Asthma | 6.7 | Military transfer orders |
| 5 | 7 | 29 | M | White | None | 3.3 | Military transfer orders |
| 6 | 6 | 34 | M | White | None | 19.9 | Participant request, adverse event |
| 7 | 6 | 45 | M | Black | Seasonal allergies<br>Allergic to shellfish<br>Eczema<br>Asthma | 20.7 | Participant request, adverse event |
| 8 | 9 | 22 | M | White | None | 96.6 | - |
| 9 | 5 | 26 | M | Asian | Seasonal allergies | 6.4 | Medically advised to stop due to adverse event |
| 10 | 5 | 24 | M | White | Anaphylaxis to raisins | 15.3 | Military transfer orders |
| 11 | 7 | 46 | M | White | Seasonal allergies | <2.0 | Two feedings skipped (medical hold, schedule conflict) |
| 12 | 9 | 27 | F | White | Urticaria to acetaminophen-hydrocodone<br>Bee allergy<br>Asthma | 26.0 | - |
| 13 | 9 | 26 | F | Multi-racial | Seasonal allergies<br>Angioedema to shellfish | 61.1 | - |
| 14 | 9 | 41 | M | White | Seasonal allergies<br>Asthma | 77.2 | - |
| 15 | 9 | 28 | F | White | Urticaria to amoxicillin-clavulanic acid | 6.7 | - |

### Progressive exposures to *Lu. longipalpis* alter the bite site rash phenotype

We characterized the phenotype and symptoms of rashes induced by uninfected *Lu. longipalpis* bites across multiple sand fly exposures. For immediate bite site reactions, 85% to 100% of participants exhibited erythema across all exposures (Fig. 2a, b), consistent with the potent vasodilatory activity of the *Lu. longipalpis* salivary protein maxadilan.^12^ Petechiae decreased with successive exposures, with only one of six participants exhibiting petechiae at the final exposure. Immediate induration was initially absent but increased to 50% of participants by the final exposure.

**Fig 2.**
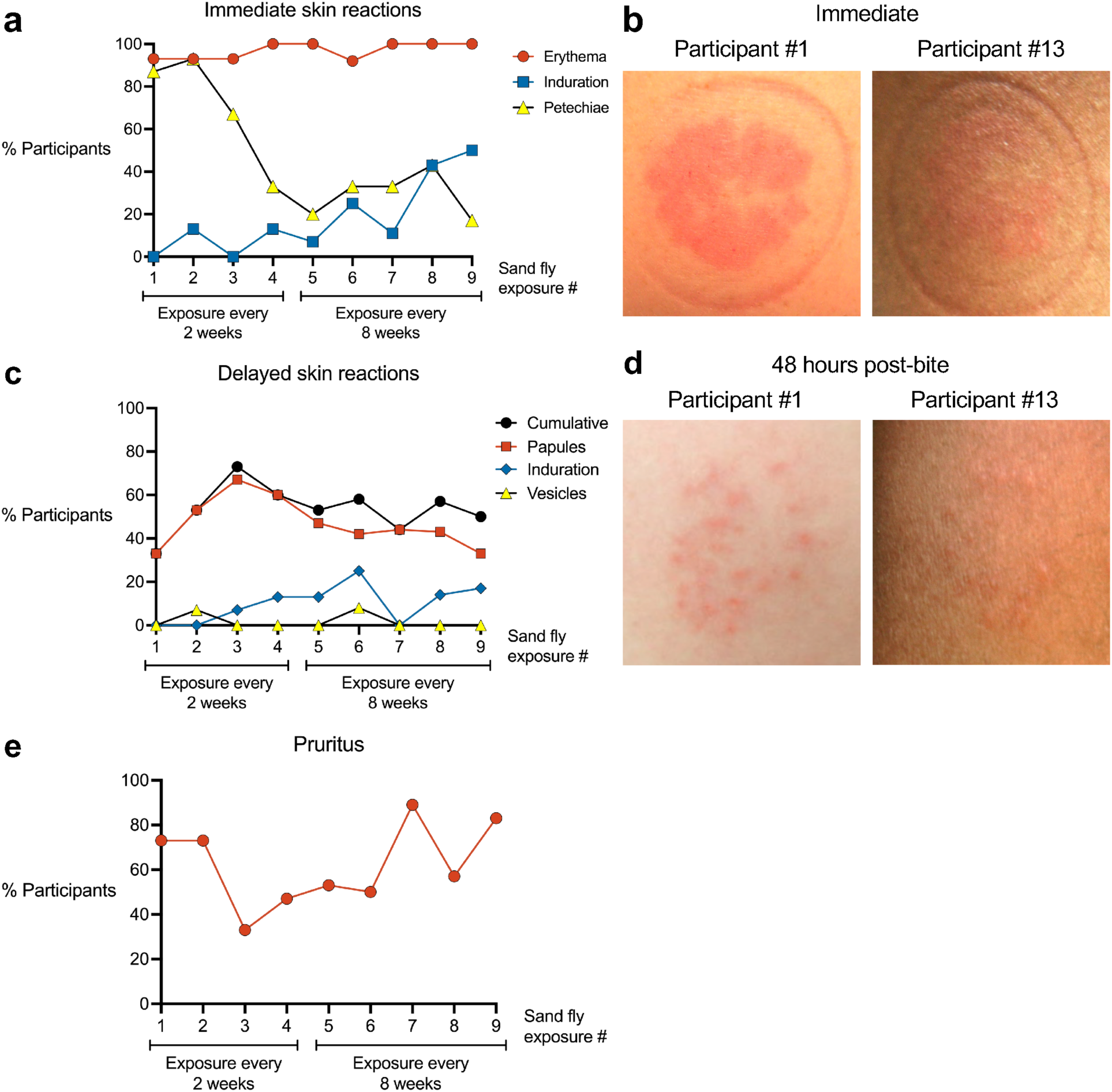
Characteristics of rashes at the sites of uninfected *Lutzomyia longipalpis* bites. **a** Skin reactions observed within the first 10 minutes after completion of each sand fly exposure. **b** Representative photographs from two participants immediately after Exposure #9. **c** Delayed skin reactions at the bite site that developed one or more days after completion of each sand fly exposure. **d** Representative photographs from two participants 48 hours after Exposure #9. **e** Pruritus at the bite site.

We next evaluated delayed bite site reactions that developed hours to one week after each sand fly exposure (Fig. 2c, d). Papules were the most frequent delayed rash phenotype (Fig. 2c), increasing from 33% to 67% of participants over the first three exposures before declining to 33% by the final exposure. Induration and vesicles appeared inconsistently in a maximum of 23% and 8% of participants, respectively. Localized pruritus at the bite site (either delayed or immediate) was initially reported by 73% of participants and increased to 83% at the final exposure (Fig. 2e). Across all exposures, the median pruritus duration was 2 days (range 30 minutes to 3 months).

Interestingly, for 8 of 15 participants, sand fly exposure triggered the appearance of a rash at a previous bite site on the contralateral arm, which had been exposed weeks before (median 6.5 weeks prior, range 2 to 11 weeks). This phenomenon, termed distal bite site reactivation, manifested approximately one week after the most recent sand fly exposure (median 4.5 exposures, range 2 to 9).

### *Lu. longipalpis* bites induce a delayed-type hypersensitivity skin response after multiple sand fly exposures

Forty-eight hours after the final sand fly exposure, two participants (#1 and #13) who completed all nine exposures volunteered for skin biopsies from a bite site and from skin free of visible inflammation on the contralateral arm as a negative control. Both participants exhibited multiple erythematous papules at their exposure site, characteristic of a DTH response (Fig. 2d).

Histology from Participant #1 showed spongiosis and a perivascular mononuclear (lymphocytic and histiocytic) infiltrate with occasional eosinophils (Fig. 3a). Participant #13 exhibited a diffuse, mixed inflammatory infiltrate with both neutrophils and mononuclear cells present in the upper dermis, distributed throughout the perivascular, interstitial, and perieccrine areas (Fig. 3b).

**Fig 3.**
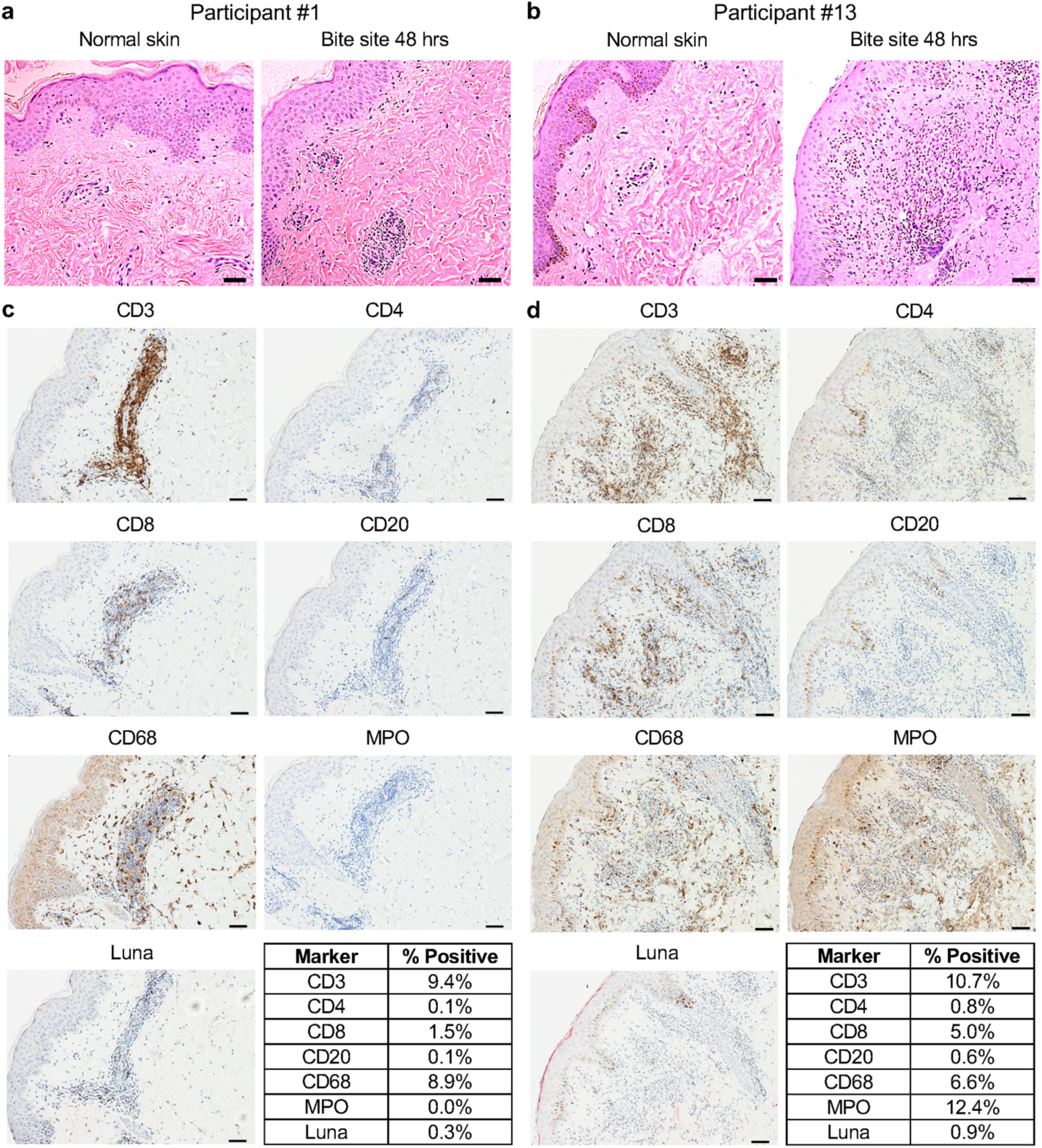
Repeated exposures to uninfected *Lu. longipalpis* bites induce a delayed-type hypersensitivity response at the bite site. Skin punch biopsies were collected from bite site skin and from normal appearing skin on the contralateral arm from two participants 48 hours after exposure #9. **a, b** Hematoxylin and eosin stains. **c, d** Immunohistochemistry (IHC) of the indicated markers at the bite site. Tables show the percentage of cells that stained positive for each marker based on analysis with ImageJ. Scale bar is 50 μm for all images.

Immunohistochemistry (IHC) revealed that the infiltrate in both participants was largely composed of T cells (CD3) and macrophages (CD68), while B cells (CD20) and eosinophils (Luna) were rare or absent, respectively (Fig. 3c, d). Interestingly, the CD4 signal was minimal (Participant #13) to absent (Participant #1), while the CD8 signal overlapped with only a fraction of the CD3 staining pattern. Thus, the CD4 and CD8 signals do not fully account for the strong CD3 signal observed in both individuals. While Participant #1 had no detectable neutrophils by myeloperoxidase (MPO) staining (Fig. 3c), Participant #13 showed neutrophil infiltration throughout the dermis (Fig. 3d). These data highlight macrophages and T cells as the predominant immune cells driving the DTH response to *Lu. longipalpis* bites in these two individuals. The IHC data also suggest that CD4^−^CD8^−^ T cells (termed double negative or DN T cells) may play a role in the DTH response to sand fly bites. The low frequency of CD4+ T cells suggests these reactions represent what we have defined as a “non-classical” DTH response where non-CD4+ T cell subsets predominate, similarly to what has been described in other skin pathologies.^13^

Quantitative RT-PCR of the skin biopsies elucidated the cutaneous TH cytokine profile at the bite site (Fig. S1). Participant #1 exhibited a mixed type 1/type 2 response with similar levels of IFN-γ and IL-13, and only minor contributions from IL-12 and IL-5. In contrast, Participant #13 showed a dominant type 1 response driven by high expression of IFN-γ, no detectable IL-13, and comparatively low expression of IL-12 and IL-5.

### *Lu. longipalpis* bites induce cellular interferon gamma responses to the sand fly salivary proteins LJL143 and LJM19

In preclinical models, distinct sand fly salivary proteins were identified as immunogenic, inducing a strong IFN-γ response which correlated with protection against leishmaniasis.^14–18^ After the second or fourth sand fly exposure, we collected peripheral blood mononuclear cells (PBMCs) from our study participants to screen the thirteen most abundant proteins in *Lu. longipalpis* saliva for their ability to stimulate IFN-γ production (Fig. 4a, Table S1). All recombinant proteins were verified to be endotoxin-free (<20 EU/ml). LJL143 and LJM19 were the only salivary proteins that stimulated IFN-γ production at a level comparable to salivary gland extract (SGE), a standard for immunogenicity. Stimulation with any of the other eleven *Lu. longipalpis* salivary proteins resulted in significantly less IFN-γ production (all *p* < 0.01). As a negative control, no IFN-γ production was elicited by SGE treatment of PBMCs from *Lu. longipalpis*-unexposed individuals obtained from the NIH Blood Bank (Fig. 4b). These results suggest that repeated exposure of humans to *Lu. longipalpis* bites leads to the development of a robust cellular IFN-γ response to the sand fly salivary proteins LJM19 and LJL143.

**Fig 4.**
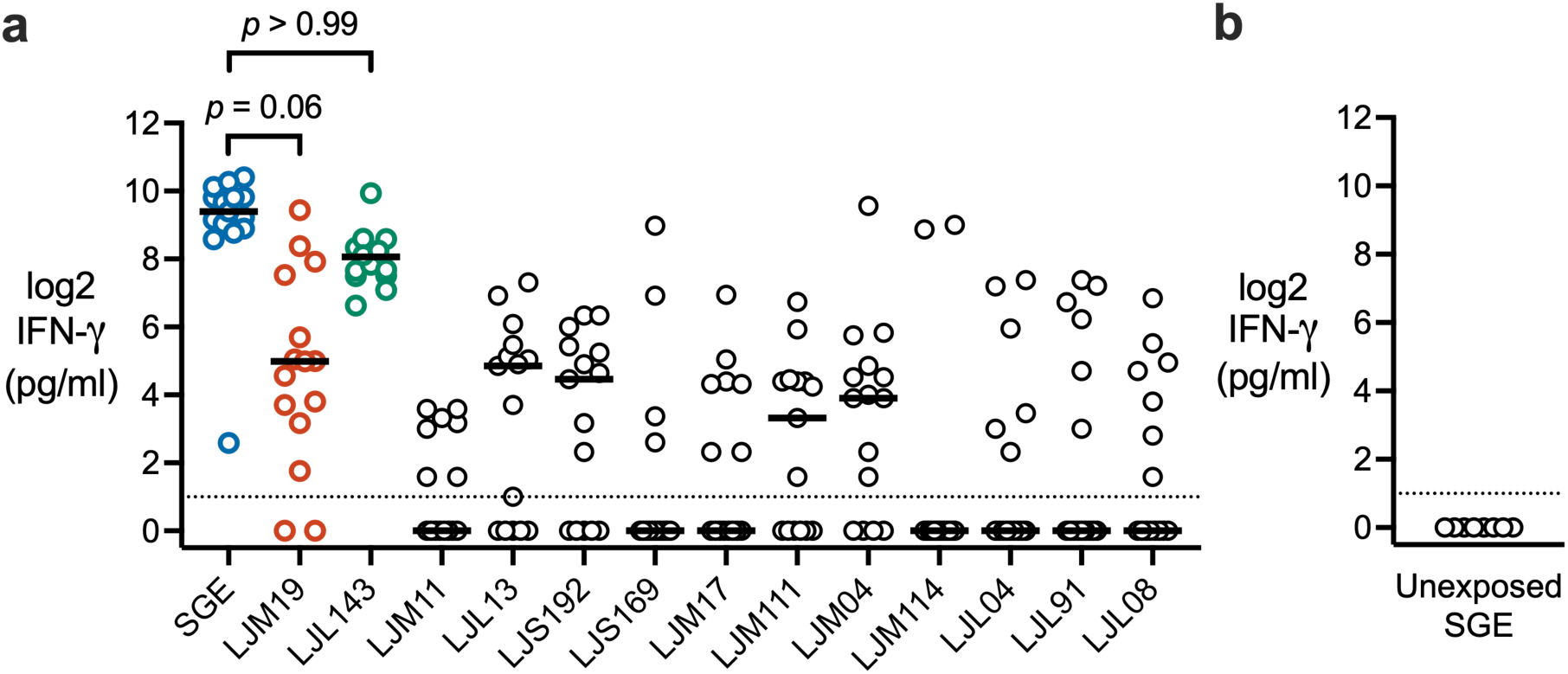
Recombinant salivary proteins stimulate IFN-γ production in PBMCs from *Lu. longipalpis*-exposed volunteers. **a** PBMCs were stimulated with salivary gland extract (SGE) or the indicated recombinant *Lu. longipalpis* salivary protein (10 μg/ml) for 96 hours to measure IFN-γ levels by ELISA. PBMCs were from exposure #2 or #4 with one time point per volunteer (*n* = 15). *P*-values are shown only for comparisons with *p* > 0.05 that showed no significant difference from SGE. **b** As a negative control, IFN-γ was measured in SGE-stimulated PBMCs from healthy volunteers unexposed to *Lu. longipalpis* (*n* = 8). Dashed lines indicate limit of detection. *P*-values were calculated by Friedman test with Dunn’s multiple comparisons test.

### LJM19 and LJL143 induce a type 1-polarized cytokine response

We next sought to elucidate the broader spectrum of TH cytokines induced by LJM19 and LJL143 compared to whole saliva (SGE). To assess these cytokine responses, we used PBMCs collected after later exposures four through nine (Table S1). PBMCs were stimulated with SGE, LJM19, or LJL143, and cytokines were measured in the supernatants by multiplex bead array and normalized via background subtraction to participant-matched PBMCs stimulated with media alone. Comparing the three stimulation conditions for each cytokine, we found that LJM19 induced significantly higher levels of IL-12 and IL-4 compared to SGE (Fig. 5a). Notably the type 2 cytokine IL-13 was consistently highly expressed across all three treatments.

**Fig 5.**
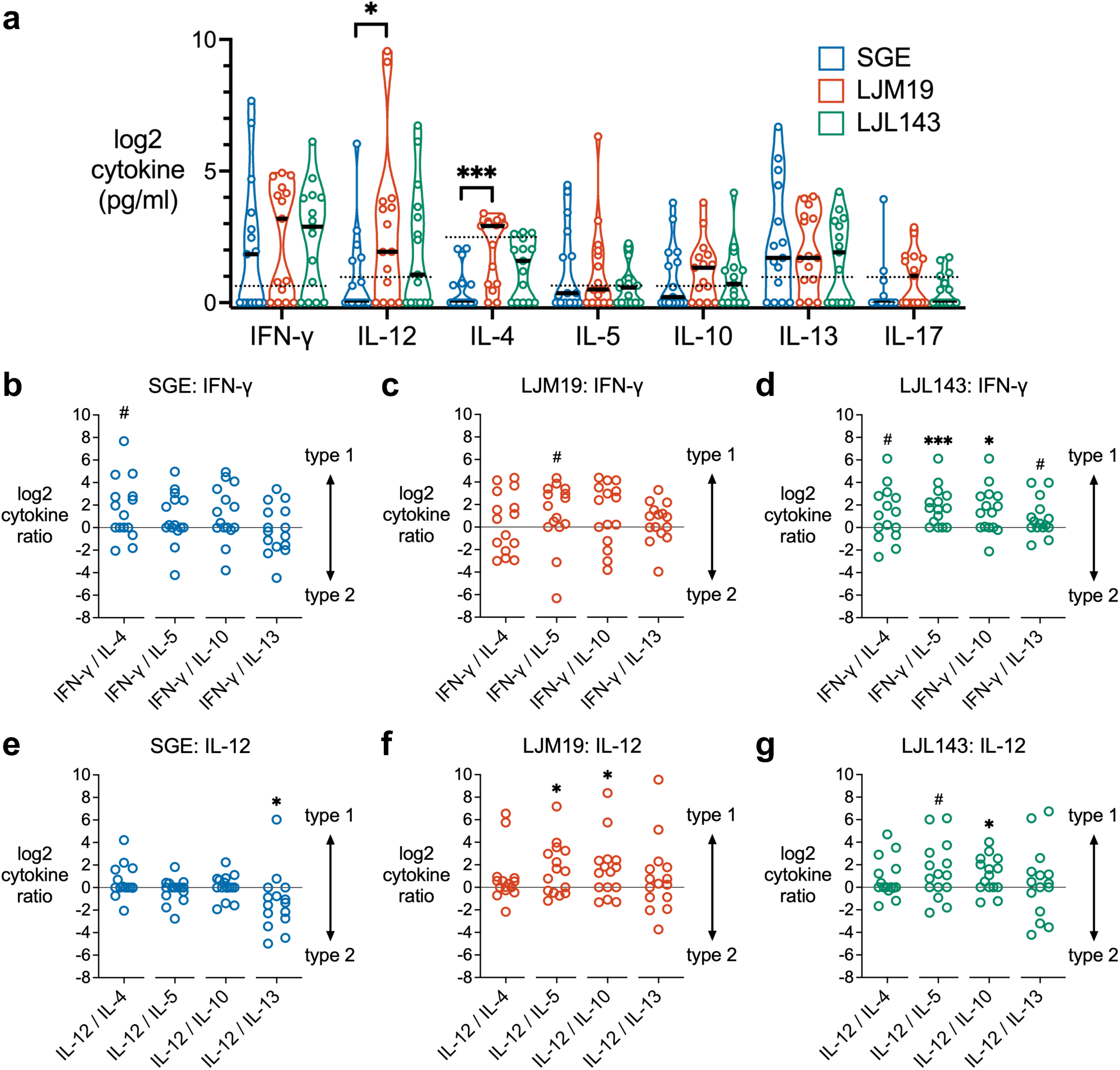
LJM19 and LJL143 induce type 1 cytokine responses in PBMCs from *Lu. longipalpis*-exposed volunteers. **a** PBMCs obtained from sand fly-exposed study participants (*n* = 15) were stimulated with SGE, LJM19, or LJL143. Cell supernatants were collected at 96 hours and cytokine concentrations were measured by multiplex bead array and normalized via background subtraction to participant-matched media-treated cells. Black bar indicates the median. Dashed lines indicate limit of detection. For each cytokine, differences between treatment groups were analyzed by Friedman test with Dunn’s test for multiple comparisons. PBMCs were from exposure #4 to #9 with one time point per participant. Ratios of type 1 to type 2 cytokines were calculated for PBMCs treated with SGE (**b, e**), LJM19 (**c, f**), or LJL143 (**d, g**) where log2 of zero represents the participant-matched, media-only baseline by design. Log ratios above 0 (solid line) indicate relative type 1 polarization while log ratios below 0 indicate relative type 2 polarization compared to media-treated controls, as analyzed by Wilcoxon signed-rank test, # *p* < 0.10, * *p* < 0.05, ** *p* < 0.01, *** *p* < 0.001.

To quantify type 1/type 2 cytokine polarization induced by SGE or each salivary protein, we calculated the ratios of the type 1 cytokines, IFN-γ and IL-12, against each type 2 cytokine (IL-4, IL-5, IL-10, and IL-13) and the type 17 cytokine, IL-17 relative to media-treated controls. We calculated cytokine ratios with either IFN-γ or IL-12 to assess both type 1 effector function and upstream regulation of TH1 lineage commitment, respectively.^19^ SGE induced a mixed type 1/type 2 response with a trend towards type 1 seen for the IFN-γ/IL-4 ratio (*p* = 0.05) (Fig. 5b). LJM19 treatment also induced a comparatively weak IFN-γ response, with a trend towards type 1 seen only for IFN-γ/IL-5 (Fig. 5c). LJL143 induced significant type 1 polarization via IFN-γ relative to IL-5 and IL-10, with a trend towards type 1 for IFN-γ versus IL-4 or IL-13 (Fig. 5d). For IL-12, SGE induced significant type 2 polarization relative to IL-13 (Fig. 5e). In contrast, LJM19-treated PBMCs exhibited significant type 1 polarization with IL-12 relative to the type 2 cytokines IL-5 and IL-10 (Fig. 5f). For LJL143, IL-12 also showed significant type 1 polarization versus IL-10 and a trend versus IL-5 (Fig. 5g). None of the treatments induced a type 17 response (Fig. S2). For each individual, the ratios were largely consistent across all type 1/type 2 cytokine pairs, except for IL-13, which was consistently expressed at a higher level than the other type 2 cytokines (Fig. S3). In summary, LJL143 and LJM19 induce a type 1-polarized response in PBMCs from individuals with multiple *Lu. longipalpis* exposures. Notably, this type 1 response is observed primarily through IFN-γ for LJL143 and through IL-12 for LJM19 and is accompanied by high IL-13 expression.

### LJM19 and LJL143 induce innate inflammatory and antiviral cytokines

We investigated the effects of SGE, LJM19, and LJL143 on other functional classes of cytokines. No differences in chemokine concentration were observed between the three treatment groups (Fig. 6a). Compared to SGE, both LJM19 and LJL143 induced significantly higher levels of interferon alpha (IFN-α), an antiviral and immunomodulatory cytokine, and IL-1β, an inflammatory cytokine (Fig. 6b). Additionally, LJM19 induced higher levels of IL-6 (Fig. 6b), an inflammatory cytokine, and IL-7 (Fig. 6c), which promotes lymphocyte proliferation and maintenance in peripheral tissues.^20^

**Fig 6.**
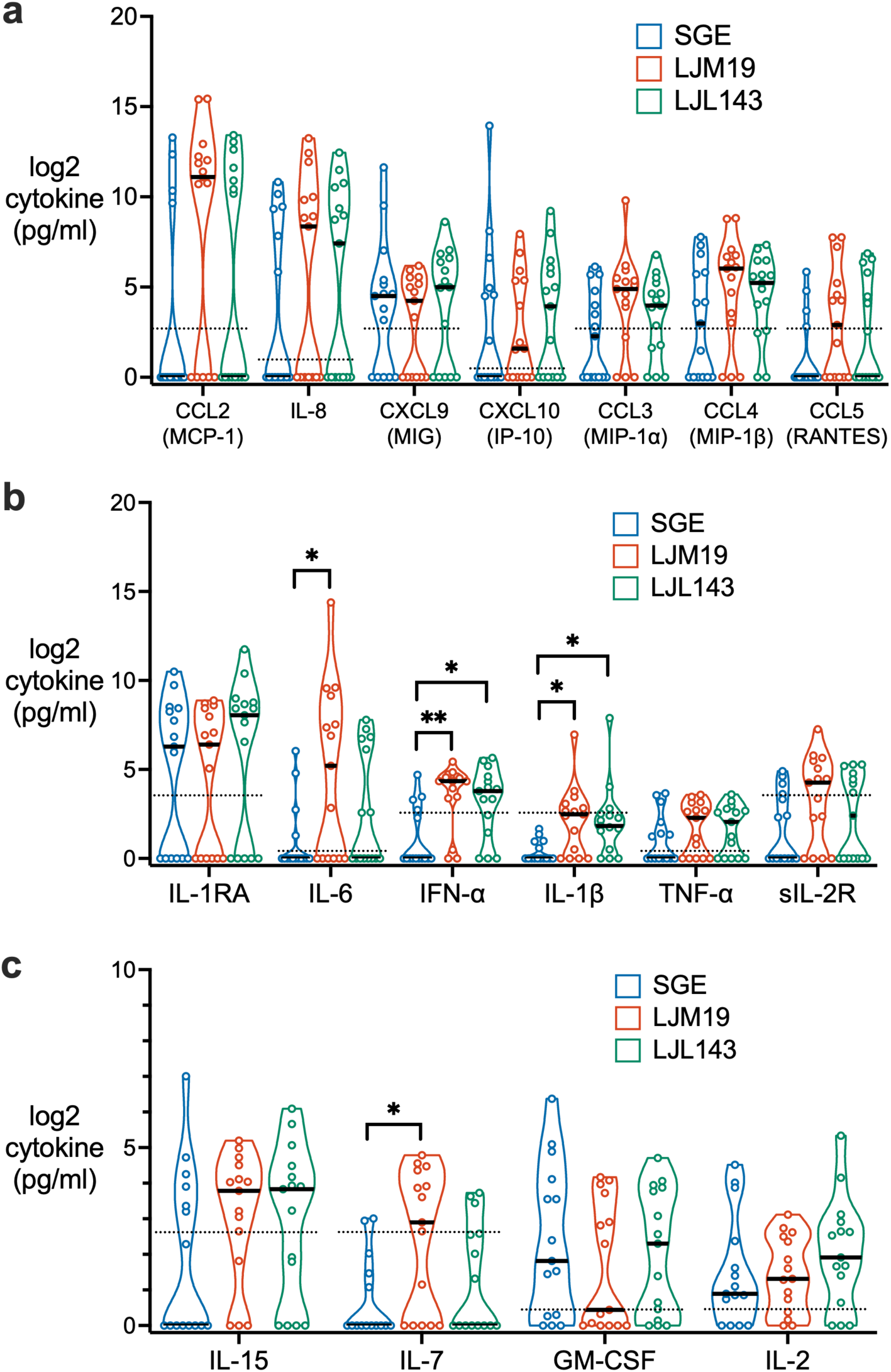
Differences in PBMC cytokine profiles induced by LJM19 or LJL143 compared to SGE. PBMCs obtained from *Lu. longipalpis*-exposed study participants (*n* = 15) were stimulated with SGE, LJM19, or LJL143. Cell supernatants were collected at 96 hours and cytokine concentrations were measured by multiplex bead array for chemokines (**a**), inflammatory cytokines (**b**), and cytokines promoting cell survival, activation, and proliferation (**c**). PBMCs used were from exposure #4 to #9 with one time point per participant. Black bar indicates the median. Dashed lines indicate limit of detection. For each cytokine, differences between treatment groups were analyzed by Friedman test with Dunn’s test for multiple comparisons, * p < 0.05, ** p < 0.01.

### LJM19 and LJL143 enhance the ability of PBMCs to stimulate macrophage killing of intracellular *Leishmania* parasites

Considering the type 1 cytokine profiles induced by LJM19 and LJL143, we tested the hypothesis that treatment with these proteins enhances the ability of PBMCs to stimulate the killing of intracellular parasites by *Leishmania*-infected macrophages. We compared PBMCs from *Lu. longipalpis*-naïve individuals from the NIH Blood Bank to PBMCs from sand fly-exposed volunteers. For the latter group, we used PBMCs collected after exposures seven through nine from a subset of four participants to assess protection against *Leishmania* in heavily sand fly-exposed individuals (Table S1). Monocyte-derived macrophages were infected with *Leishmania infantum* for eight hours, then co-cultured with autologous PBMCs stimulated with the sand fly salivary proteins LJM19 or LJL143. We sought to capture the effects of LJM19 and LJL143 on early macrophage infection by *Leishmania*, because this is the time window when the proteins would begin to enhance parasite killing as candidate vaccine antigens and adjuvants. Phytohemagglutinin (PHA), a T cell mitogen, served as a positive control (Fig. 7a).

**Fig 7.**
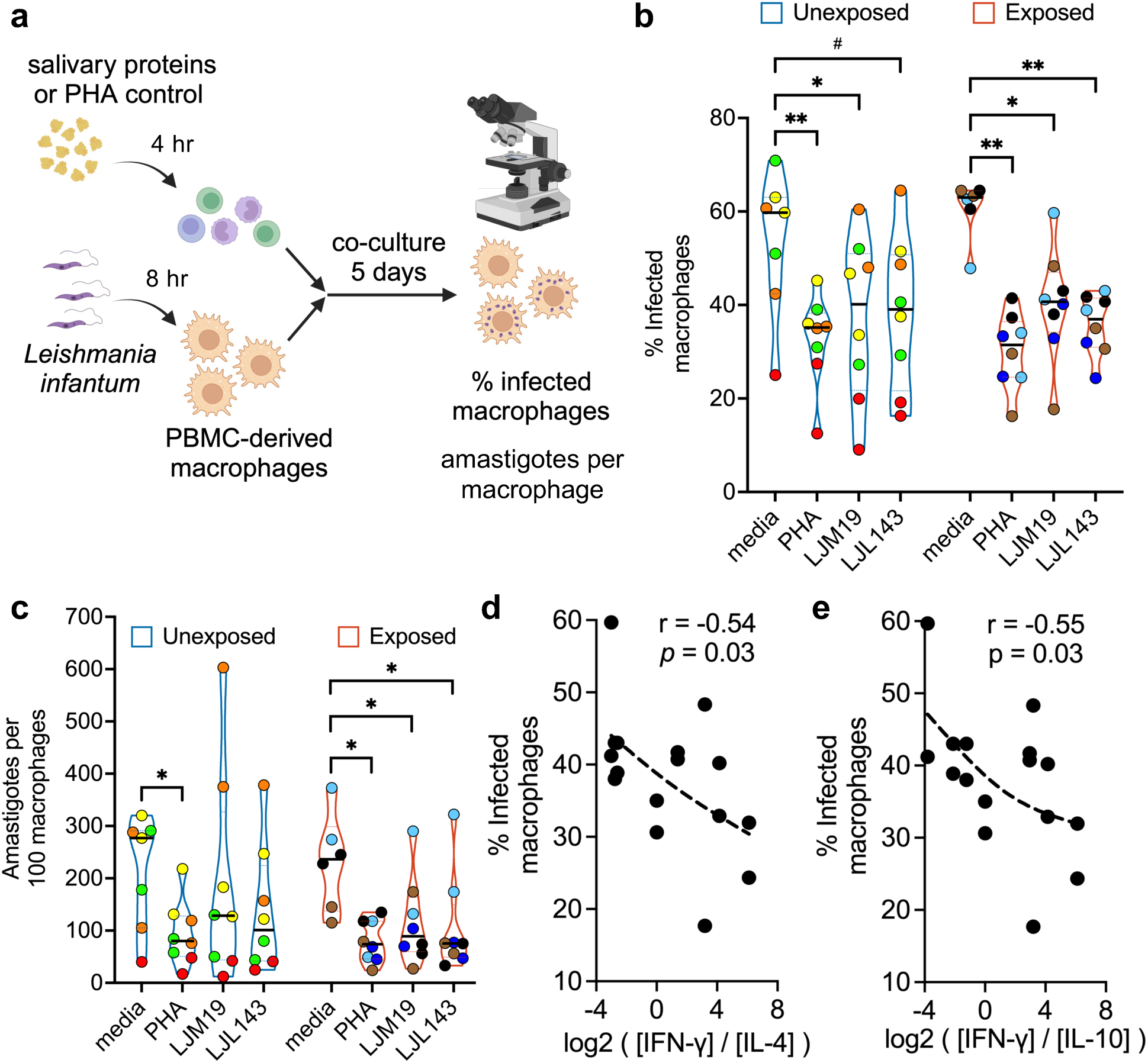
Stimulation of PBMCs with LJM19 or LJL143 enhances killing of *Leishmania* by macrophages. PBMC-derived macrophages from participants unexposed (*n* = 4, blue border) or exposed (*n* = 4, red border) to *Lu. longipalpis* were infected with *Leishmania infantum*, then co-cultured with autologous PBMCs that had been stimulated according to the conditions shown. After 5 days of co-culture, the percentage of infected macrophages was quantified by manual counting of Giemsa-stained cells by light microscopy. PBMCs were from exposure #7 to #9 with one time point per participant. **a** Experimental scheme. **b, c** Each volunteer’s batch of PBMCs was divided and run in two technical replicates; identical colors denote replicates from the same individual. The percentage of infected macrophages (**b**) and the number of amastigotes per 100 macrophages (**c**) were measured. Bar indicates the median. Differences between treatment groups were analyzed via a mixed effects model with Holm-Šídák’s test for multiple comparisons, # *p* < 0.10, * *p* < 0.05, and ** *p* < 0.01. **d, e** Spearman correlation between the IFN-γ/IL-4 ratio (**d**) or IFN-γ/IL-10 ratio (**e**) as calculated in Figure 5 and the percentage of infected macrophages for LJM19- and LJL143-treated samples. Dashed line in **d, e** is the trend line (smoothing spline). Schematic in **a** created in BioRender.com.

For *Lu. longipalpis*-exposed participants, PBMC stimulation with either LJM19 or LJL143 led to a significant reduction in the percentage of infected macrophages (Fig. 7b) and the number of amastigotes per macrophage (Fig. 7c). These reductions were comparable in magnitude to that observed with PHA stimulation. Surprisingly, in PBMCs from *Lu. longipalpis*-unexposed individuals, stimulation with LJM19 alone also induced a significant decrease in the percentage of infected macrophages, while stimulation with LJL143 showed a trend towards a decrease (Fig. 7b). In contrast, the corresponding decreases in amastigotes per macrophage did not meet the significance threshold (Fig. 7c). When comparing unexposed to exposed individuals, there were no significant differences in the percentage of infected macrophages (*p* = 0.70 for LJM19; *p* = 0.68 for LJL143) or the number of amastigotes per macrophage (*p* = 0.79 for LJM19; *p* = 0.75 for LJL143). These data suggest that LJM19 and potentially also LJL143 possess intrinsic innate immune activating properties that promote *Leishmania* killing. In contrast, sand fly exposure induced a more consistent adaptive response potentially further enhancing parasite killing for both salivary proteins, particularly by reducing amastigote load (Fig. 7c).

Cross-referencing these macrophage infection data with our earlier type 1/type 2 cytokine ratio results from sand fly-exposed individuals revealed a significant negative correlation between the percentage of infected macrophages and the ratios of IFN-γ/IL-4 and IFN-γ/IL-10 (Fig. 7d, e), while no correlation was seen with the IFN-γ/IL-5 or IFN-γ/IL-13 ratios (Fig. S4). Similarly, no significant correlation was seen with IFN-γ alone (Fig. S4), indicating that IFN-γ is not the sole determinant of the parasite killing effect mediated by LJM19 and LJL143, but rather that the killing effect depends on the type 1/type 2 cytokine balance.

## DISCUSSION

This study provides the most comprehensive characterization to date of how human skin and systemic immune responses evolve with longitudinal exposure to the bites of *Lutzomyia longipalpis* sand flies. Our cohort was larger than the only prior study to perform controlled human challenge with *Lutzomyia*,^4^ and had a participant retention rate comparable to a recent human challenge study with *Phlebotomus duboscqi*.^10^ All participants completed at least five exposures, which is sufficient to develop adaptive immunity against sand fly saliva and surpasses the prior benchmark of four exposures from the previous *Lutzomyia* challenge study.^4^ Moreover, several participants in our cohort completed up to nine exposures over the course of a year.

Stable DTH reactions to *Lu. longipalpis* bites developed and were maintained throughout the study, similar to those seen previously against *Phlebotomus duboscqi* bites.^5,10^ In preclinical models, saliva-specific type 1 DTH responses are closely correlated with immune protection against *Leishmania*.^16,17^ In our study, skin biopsies of the bite site from two participants who completed all nine sand fly exposures exhibited variable DTH reactions that were either type 1 or exhibited a mixed type 1/type 2 response. In both volunteers, the DTH response consisted predominantly of T cells and macrophages, with neutrophils present in one participant. Notably, both biopsies showed a relative absence of CD4+ T cells, and were instead dominated by CD3+ CD4– CD8– double negative (DN) T cells, consistent with a non-classical DTH phenotype. While we cannot formally exclude the possibility of incomplete detection of CD4 by IHC, our report of DN T cells corroborates prior findings in a cohort from Mali where three of six individuals had a DTH response to sand fly bites characterized by infiltration with DN T cells, while CD4+ and CD8+ T cells were absent.^5^ Although the role of DN T cells in responses to sand fly bites remains incompletely defined, expanded DN T cell populations have been observed in human CL lesions due to *L. braziliensis* and exhibit a highly activated phenotype.^21,22^ These DN T cells express an αβ T cell receptor (TCR) and show high expression of IFN-γ and CD69, a tissue residence marker.^21^ In mice, DN T cells are necessary for primary and secondary immunity to *L. major*.^23^ Further studies utilizing RNA-seq and flow cytometry are needed to better define the immune cell subsets infiltrating the sand fly bite site in humans, to elucidate the dominant TH polarization profile at the sand fly bite site, and to determine whether DN T cells are a consistent and prominent feature of the DTH response that influences immunity to *Leishmania* parasites.

In over half of participants, *Lu. longipalpis* bites triggered the reappearance of a rash at a previously bitten site on the contralateral arm. This phenomenon, known as distal bite site reactivation, was also reported after uninfected *Phlebotomus duboscqi* bites^10^ and during early studies of human skin reactions to the bites of *Phlebotomus papatasi*.^24^ We speculate that this reactivation reflects activation of salivary protein-specific skin resident memory T cells that have seeded a prior bite site and respond to circulating salivary antigens from new sand fly bites.^25,26^ The ability of vector bites to impact systemic immunity was demonstrated in humanized mice exposed to uninfected *Aedes aegypti* bites.^27^ These mice exhibited significant changes in blood cytokine levels and immune cell composition in both blood and skin up to seven days after mosquito exposure. This suggests the intriguing possibility that local cutaneous responses to vector bites may trigger systemic signals that promote a pathogen-resistant state throughout the skin, similar to the organism-wide coordination of antiviral immunity.^28,29^

Using PBMCs, our screen identified LJM19 and LJL143 as the two *Lu. longipalpis* salivary proteins that stimulate the highest IFN-γ production in PBMCs from sand fly exposed participants, confirming their immunogenicity in humans. Unexpectedly, treatment of PBMCs from *Lu. longipalpis*-unexposed individuals with LJM19 or LJL143 reduced the percentage of *Leishmania*-infected macrophages in co-culture. These findings in human PBMCs corroborate data from preclinical models demonstrating that LJM19 and LJL143 have intrinsic adjuvant-like properties that enhance the immunogenicity of parasite-derived vaccine antigens.^30,31^ LJM19 (also named SALO) is an 11 kDa protein that inhibits the classical complement pathway and has no structural similarity to human proteins.^32,33^ LJM19 was previously identified as a salivary protein vaccine candidate that protects against both cutaneous and visceral leishmaniasis in preclinical models.^15,18,31^ LJL143 (also named Lufaxin) is an inhibitor of coagulation factor Xa and the alternative pathway of complement.^34,35^ In dogs, LJL143 elicits strong DTH and IFN-γ responses in the skin and blood following uninfected sand fly challenge.^14^ LJL143 exhibits adjuvant-like activity in mice, where priming immunized animals with unadjuvanted LJL143 induced higher CD4+ T cell proliferative responses to *Leishmania* antigens in a virus-like particle (VLP) vaccine compared to mice that did not receive LJL143.^30^ Together, these findings suggest that the innate immune-activating properties of LJM19 and LJL143 enhance their efficacy as salivary protein vaccine candidates against leishmaniasis. Identifying the innate immune receptors and cell subsets that mediate the adjuvant-like effects of LJM19 and LJL143 remains an area for future research.

Beyond their innate immune-activating effects, repeated sand fly exposure enhanced the ability of both LJM19 and LJL143 to reduce amastigote burden in macrophages. Furthermore, the reduction in the percentage of infected macrophages correlated with type 1 cytokine polarization. In participants exposed to *Lu. longipalpis*, treatment of PBMCs with SGE induced a mixed type 1/type 2 response, while LJM19 and LJL143 both induced type 1-polarized responses. Taken together, these findings are consistent with a model where repeated sand fly exposure leads to the development of type 1 adaptive immunity specific to LJM19 and LJL143. LJM19 induced high expression of the type 1 cytokine IL-12, whereas LJL143 induced high expression of both IFN-γ and IL-12. Classically, IL-12 is secreted by professional antigen presenting cells and regulates TH1 lineage commitment, leading to IFN-γ production by T cells.^19^ However, as seen with LJM19, strong induction of IL-4 can suppress IFN-γ despite high levels of IL-12.^36^ This co-expression of IL-12 and IL-4 mirrors the cytokine profile seen in LJM19-immunized hamsters protected against *L. donovani* challenge.^31^ These patterns underscore that salivary protein-induced immunity does not conform to a binary type 1/type 2 model, but instead reflects a context-dependent balance of opposing regulatory signals.

Of the type 2 cytokines, IL-13 was often expressed at levels equal to or higher than IFN-γ. IL-13 promotes *Leishmania* infection and exacerbates immunopathology,^37–39^ in part by inhibiting expression of IL12Rβ2 which transduces critical signals for type 1 differentiation.^40,41^ Polymorphisms at the *IL13* locus in humans have also been associated with susceptibility to CL caused by *Leishmania guyanensis*.^42^ In a naturally exposed population in Mali, IL-13 was highly expressed in PBMCs stimulated with SGE from *P. duboscqi*, but only at low levels in the skin after uninfected bite challenge.^5^ Further studies will be needed to determine whether the high IL-13 expression induced by LJM19 and LJL143 reflects an intrinsic property of these proteins or a counterregulatory response to type 1 polarization.

In addition to type 1 and type 2 effector cytokines, we found that LJM19 induces IL-6 and IL-7, while both LJM19 and LJL143 induce IL-1β and the type I interferon, IFN-α. This contrasts with the reported downregulation of IL-6 and IL-1β by PpSP32, another sand fly salivary protein.^43^ To our knowledge, this is the first study to report altered expression of IL-7 and IFN-α in response to specific sand fly salivary proteins. IL-6 exerts pleiotropic immune effects, including activation of the acute phase response, granulopoiesis, B cell proliferation, and CD8+ T effector cell development.^44^ In mice, IL-6 has been shown to facilitate resistance to *L. major*^45^ and *L. donovani*^46^, in the latter case by inhibiting the proliferation of IL-10-expressing CD4+ T cells. IL-7 supports the survival, proliferation, and maintenance of T cells in peripheral tissues,^20^ including skin resident memory T cells,^47^ which are critical for sustained protection against *Leishmania*.^48^ While IL-1β has been reported to protect against *L. amazonensis*,^49^ most studies suggest IL-1β exacerbates parasite dissemination and immunopathology.^50–54^ Similarly, IFN-α likely benefits the parasite by antagonizing IFN-γ responses and suppressing the development of *Leishmania*-specific T cells,^55,56^ although the timing of IFN-α signaling relative to *Leishmania* infection can shift the balance between protection and susceptibility.^57^

Our study has several limitations. The cohort is demographically homogenous (80% male, 73% White), limiting generalizability. Due to limited PBMC recovery at some blood collections, each assay used PBMCs from more than one time point. While we cannot exclude the possibility that participants were previously exposed to local *Lutzomyia* species at low prevalence,^58^ all participants and blood bank samples were screened and found to be negative for IgG against *Lu. longipalpis* SGE. Due to their differentiation in GM-CSF, the PBMC-derived macrophages in the parasite killing assay were likely polarized to the inflammatory M1 phenotype,^59^ which would enhance their baseline ability to kill intracellular pathogens. Nevertheless, because all macrophages were differentiated under identical conditions, comparisons across treatments remain internally consistent. Finally, most of our findings are based on PBMC responses and need reinforcement through studies of immune cells in the skin.

In summary, we report that humans exposed to *Lu. longipalpis* generate robust innate and adaptive cellular immune responses to the sand fly salivary proteins LJM19 and LJL143. These proteins exhibit adjuvant-like activity, induce type 1 cytokine polarization, and enhance the ability of macrophages to kill *Leishmania*. Our findings suggest a model where innate immune signals induced by LJM19 or LJL143 work in concert with adaptive type 1 immunity in *Lutzomyia*-exposed individuals to bolster the anti-*Leishmania* response. This study highlights the potential of leveraging anti-saliva immunity in humans to protect against vector borne pathogens. Furthermore, we posit that chronic immune stimulation by vector bites is underappreciated as an environmental exposure that profoundly reshapes human immune responses in endemic populations. Our work underscores the importance of human vector challenge studies as an approach to investigate how repeated exposure to vector salivary antigens impacts human immunity, a daily occurrence for the approximately 350 million people who live in regions where sand flies and other disease vectors are prevalent.

## METHODS

### Study approval

The Institutional Review Boards of Walter Reed Army Medical Center, Walter Reed National Military Medical Center (protocol number WR355023), the National Institute of Allergy and Infectious Diseases (NIAID), and the Uniformed Services University of the Health Sciences (USUHS) approved this study. All human subjects research was conducted in accordance with the principles of the Declaration of Helsinki and complies with all relevant ethical regulations. Participants provided written informed consent prior to study participation, including consent for the use of their photographs. The study was registered on www.clinicaltrials.gov as NCT01289977.

### Study population

The screening cohort and enrollment criteria have been published previously for a related study on human immunity to the bites of *Phlebotomus duboscqi*, a sand fly vector of cutaneous leishmaniasis predominantly found in Africa,^10^ however there was no overlap in the enrollment cohort of the prior study and the current one. In this single site study conducted at Walter Reed Army Medical Center (WRAMC), 68 healthy individuals were screened and 15 were enrolled in the *Lu. longipalpis* study cohort. Study participants provided self-identified demographic information, including race and sex, using options defined by the investigator. A medical history and physical examination were obtained and included a review of allergies and travel history. The inclusion criteria were healthy military healthcare beneficiaries between 18 and 50 years old who were willing to remain in the local area for the next 12 months and participate in all study procedures. Exclusion criteria included a history of travel for more than 30 consecutive days to a geographic area where *Lutzomyia longipalpis* is present, positivity by screening ELISA to IgG that bind *Lu. longipalpis* salivary gland extract (SGE), elevated serum IgE >144 kU/L, pregnancy, history of chronic medical illness, large skin reactions to insect bites, problems with prior phlebotomy, or use of medications that may interfere with immune responses.

### Human controlled exposure to uninfected laboratory-reared *Lutzomyia longipalpis*

The colony of *Lu. longipalpis* sand flies used for this study was originally field collected in 2004 in Jacobina, Brazil. For the clinical study, *Lu. longipalpis* were reared in a pathogen-free insectary at the Laboratory of Malaria and Vector Research (LMVR), NIAID, and were maintained as a closed colony. For each participant, 10 female *Lu. longipalpis* sand flies were starved overnight, loaded into a feeding chamber, then transported to WRAMC on the day of the exposure. Each feeding chamber is composed of a sealed Plexiglass capsule with a fine mesh surface (Precision Plastics, Inc.). The feeding chamber was secured to the upper arm of each participant with the mesh side contacting the skin, allowing the sand flies to feed through the mesh. Each sand fly exposure lasted 20 minutes, during which the feeding chamber was lightly covered with fabric to promote a dark feeding environment. Areas of skin with tattoos were avoided for feeding sites. At the end of each exposure, all sand flies were accounted for and examined by microscopy to assess the number of flies that had taken a blood meal.

Bite site skin reactions were observed by study physicians for 10 minutes immediately following the end of the sand fly exposure. Any participants with skin reactions deemed to be large or potentially allergic were observed for a longer period. Bite site photographs were taken and the physical appearance of the bite site rash and any associated symptoms were recorded. Participants were counseled to refrain from using antihistamines or topical steroids for bite site symptoms until consultation with study physicians. Sand fly exposures were performed once every two weeks for exposures #1 through #4, then once every eight weeks for exposures #5 through #9 (Fig. 1). Feeding sites were alternated between different arms on consecutive visits. Blood was collected from participants 7 ± 3 days following each sand fly exposure, at which time the bite site was reassessed for delayed skin reactions.

### Skin histology and immunohistochemistry

Forty-eight hours after their final exposure (#9) to *Lu. longipalpis*, two study participants (#1 and #13) consented to have a 3 mm skin punch biopsy (Miltex sterile skin punch biopsy tool) taken at a bite site from the most recent exposure and a 2 mm punch biopsy from normal appearing skin free of visible inflammation on the contralateral arm as a negative control. Each biopsy was bisected, with one half stored in 10% buffered formalin for histology and the other half stored in RNAlater (Ambion) for quantitative (real time) RT-PCR.

Embedding of the biopsies as formalin-fixed paraffin-embedded (FFPE) blocks and histological staining was performed by Histoserv (Germantown, Maryland). Five micron-thick tissue sections were stained with hematoxylin and eosin and evaluated by light microscopy. For immunohistochemistry (IHC), primary antibodies against the following targets were used at the dilutions listed: CD3 at 1:100 (Dako #A0452), CD4 at 1:80 (Dako #M7310), CD8 at 1:75 (Dako #M7103) at 1:75, CD20 at 1:300 (Dako #M0755), CD68 at 1:100 (Dako #M0814), and myeloperoxidase (MPO) at 1:400 (Dako#A0398). For secondary antibodies, biotinylated anti-mouse IgG (1:500) was used to detect primary antibodies binding CD4, CD8, CD20, CD68, and biotinylated anti-rabbit IgG (1:500) was used to detect anti-CD3. Streptavidin-horseradish peroxidase was used to visualize the protein targets. Slide photography was performed using an Olympus DP73 camera microscope BX51 with Cellsens Dimension Olympus software. The percentage of cells positive for each IHC marker was quantitated with ImageJ software.

### Quantitative RT-PCR of skin cytokines

For each biopsy portion stored in RNAlater, RNA was extracted using the RNeasy Fibrous Tissue Mini Kit (Qiagen) and treated with DNase I to remove contaminating genomic DNA. Total RNA (100 ng) was used for cDNA synthesis using the qScript cDNA Supermix (Quanta Biosciences). Absence of genomic DNA contamination was verified by PCR of total RNA. Relative quantification of IFN-γ, IL-12, IL-4, IL-5, and IL-13 was performed in a LightCycler 480 (Roche Applied Science) using the Universal ProbeLibrary system (Roche). Primers and probes were designed using ProbeFinder software (v 2.45, Roche). Relative quantification of target genes normalized to 18S rRNA was performed using the LightCycler 480 software. Cytokine gene expression from the bite site biopsies was then normalized to the skin biopsy from the contralateral arm.

### Preparation of *Lu. longipalpis* salivary gland extract

Salivary gland extract (SGE) was prepared by dissection of salivary glands from seven day old, laboratory-reared, uninfected adult female *Lu. longipalpis*. Glands were homogenized by ultrasonication with a Branson Sonifier 450 for three 30 second cycles then clarified by centrifugation at 10,000 *xg* for 3 min at 4 ^O^C. Supernatant extracts were collected and stored at −80 ^O^C until use.

### Cloning and expression of *Lu. longipalpis* salivary proteins

DNA of the most abundant salivary molecules from *Lu. longipalpis* was amplified by polymerase chain reaction (PCR) using a forward primer derived from the amino-terminal sequence immediately 3’ to the signal peptide sequence and a reverse primer encoding a hexa-histidine tag. The PCR conditions were: one hold for 5 min at 94 ^O^C, two cycles of 30 s at 94 ^O^C, 1 min at 46 ^O^C, 1 min at 72 ^O^C and 23 cycles of 30 s at 94 ^O^C, 1 min at 52 ^O^C, 1 min at 72 ^O^C and one hold of 7 min at 72 ^O^C. The PCR product was cloned into the VR2001-TOPO vector as previously described^60^ then sequenced. The VR-2001 plasmid encoding the His-tagged salivary proteins was sent to the Protein Expression Laboratory at NCI Frederick (Frederick, Maryland) for expression in HEK-293F cells. Supernatant was collected after 72 hours and concentrated from 1 L to 300 ml using a stirred ultrafiltration cell unit (Millipore) with an ultrafiltration membrane (Millipore). The volume was restored to 1 L by the addition of 500 mM NaCl and 10 mM Tris, pH 8.0. The protein was purified by HPLC (Biorad, NGC Chromatography System) using two tandem 5 ml HiTrap Chelating HP columns (GE Healthcare) charged with 0.1 M NiSO4. Protein was detected at 280 nm and eluted by an imidazole gradient. Eluted proteins were collected every minute in a 96-well microtiter plate using a BioFrac fraction collector (Biorad). Fractions corresponding to specific absorbance peaks were selected and run on a NuPage Bis-Tris 4–12% Gel (Novex) with MES running buffer under reducing conditions per the manufacturer’s instructions. Afterwards, the gel was stained with 0.025% Coomassie blue to visualize proteins. Specific fractions were selected based on molecular weight and concentrated to 1 ml using an Amicon Ultra Centrifugal Filter (Millipore). The protein sample was then injected into a G2000SW molecular sieving column (Tosoh Biosciences) via a 1 ml loop connected to the HPLC (DIONEX) with phosphate buffered saline (PBS), pH 7.2 as the buffer for further purification steps. Protein was detected by absorbance at 280 nm and the fractions were of interest were collected and pooled as described above. Protein concentration was measured with a NanoDrop ND-1000 spectrophotometer at 280 nm and calculated using the extinction coefficient of the protein. Endotoxin levels were measured with the Endosafe Portable Test System (Charles River Laboratories) and verified to be <20 EU/ml for all proteins.

### Blood collection and storage

Blood was collected from each study participant in heparinized Vacutainer tubes (BD Diagnostics). Collections were performed after exposures #2, #4, and #5 through #9 (Fig. 1). Peripheral blood mononuclear cells (PBMCs) were isolated by density gradient centrifugation using a Ficoll-Paque PLUS solution (GE Healthcare). Plasma supernatants were collected and stored at −80 ^O^C. PBMCs were counted, resuspended in fetal bovine serum (FBS) with 10% dimethyl sulfoxide (DMSO) solution, and transferred to cryovials which were slowly cooled to −80 ^O^C overnight in a Mr. Frosty freezing container (Thermo Fisher Scientific) then transferred to liquid nitrogen.

To obtain pre-exposure negative controls, PBMCs were collected from each study participant by apheresis prior to sand fly exposure. However, initial tests of these apheresed PBMCs demonstrated that they were broadly reactive to the majority of recombinant sand fly salivary proteins tested, indicating that the apheresis procedure had caused non-specific activation of these PBMC batches. Thus, we elected to use PBMCs collected from healthy volunteers at the NIH Blood Bank as *Lu. longipalpis* unexposed negative controls in our stimulation assays (Fig. 4 and Fig. 7). Blood from each donor was screened by ELISA to verify they were negative for IgG against *Lu. longipalpis* SGE. All post-exposure blood was collected via conventional venous phlebotomy.

### Peripheral blood mononuclear cell culture and stimulation

For stimulation assays, cryopreserved PBMCs were thawed quickly at 37 ^O^C, diluted in RPMI 1640 medium, then pelleted by centrifugation for 10 min at 350 *xg* at RT. PBMCs were cultured in RPMI 1640 medium supplemented with 10% AB human serum (Sigma), 1% sodium pyruvate, 1% non-essential amino acids, 1% HEPES buffer, 50 µM beta-mercaptoethanol, and 40 mg/ml penicillin/streptomycin in a 5% CO2 humidified atmosphere at 37 ^O^C overnight. Cells were counted and viability was assessed using Trypan blue (Hyclone, Thermo Fisher Scientific). PBMCs were then cultured in 96-well plates in cell culture medium at 1 x 10^6^ cells/ml in a final volume of 200 μl per well and incubated with medium alone, SGE (0.5 pairs/ml), 10 μg/ml recombinant *Lu. longipalpis* salivary proteins, or 2.5 μg/ml concavalin A (ConA). Cell supernatants were collected after 96 hours, clarified by centrifugation for 10 min at 350 *xg* at 4 ^O^C, then stored at −80 ^O^C until further use.

Within each assay, PBMCs from each individual were taken from a single time point, though the time point used varied between individuals, depending on the availability of PBMCs. The specific time points used for each assay for each individual are detailed in Table S1.

### Interferon gamma ELISA

ELISA was performed on thawed supernatants of stimulated PBMCs from exposure #2 or #4 (Table S1) using the anti-human interferon gamma ELISA kit (BD Biosciences) according to the manufacturer’s instructions. The results were interpolated from a standard curve using recombinant IFN-γ and expressed as the concentration of IFN-γ (pg/ml) minus the amount of background IFN-γ secreted from cells treated with media alone.

### Cytokine multiplex bead array

Cytokine concentrations in the supernatants of PBMCs from exposure #4 through #9 (Table S1) were measured using the Cytokine Human Magnetic 25-Plex Panel (Invitrogen) according to the manufacturer’s instructions. Results reported reflect the concentration of each secreted cytokine (pg/ml) minus the background concentration of cytokine from participant-matched cells treated with media alone.

### PBMC-macrophage co-culture *Leishmania* killing assay

Human macrophages were derived from PBMCs of exposures #7 through #9 (Table S1) in a subset of study participants (*n* = 4) who completed all nine *Lu. longipalpis* exposures or from NIH blood bank volunteers (*n* = 4). Two technical replicates were performed on PBMCs from each participant, for a total of 8 samples per sand fly exposure group. For differentiation into macrophages, PBMCs were plated in a 16-well chamber slide at 3 x 10^5^ cells per well in the presence of complete RPMI and 20 ng/ml of GM-CSF for 1 hour at 37 ^O^C with 5% CO2. Non-adherent cells were removed, then media was replaced with complete RPMI + 20 ng/ml GM-CSF. On day 5, matched donor PBMCs were thawed for use in the autologous killing assay and cultured in suspension in RPMI. On day 6, the cultured macrophages were infected with stationary phase *L*. *infantum* at a parasite:macrophage ratio of 5:1, then incubated for 8 hours at 37 ^O^C with 5% CO2. Non-internalized parasites were removed by washing. Autologous PBMCs were treated with media, phytohemagglutinin (PHA) (6.25 µg/ml), or salivary protein (10 µg/ml) for 4 hours at 37 ^O^C with 5% CO2, then added to the *Leishmania*-infected macrophages at 5 x 10^5^ PBMCs per well. After 5 days of co-culture, cells were fixed, stained with Giemsa, and assessed by light microscopy. Approximately 300 macrophages were counted per well and evaluated for the number of intracellular *Leishmania* amastigotes.

### Statistics and reproducibility

Statistical analyses were performed in GraphPad Prism (v 10.2.0). Differences were considered statistically significant for p < 0.05 (*). Due to the exploratory nature of our study, we also indicated where p > 0.05 but < 0.10 (denoted by “#”) to identify potential trends that may be biologically relevant but did not achieve the significance threshold. All statistical tests were two-sided, where applicable.

For the IFN-γ ELISA (Fig. 4), non-parametric statistical tests were used because the dataset failed both normality and lognormality tests. Friedman test with Dunn’s multiple comparisons test was used for pairwise comparisons of IFN-γ concentration between SGE (comparator) and each of the *Lu. longipalpis* recombinant salivary proteins.

For cytokine measurements by multiplex bead array (Fig. 5 and Fig. 6) and cytokine ratio analyses, cytokine concentrations were first background-subtracted using participant-matched media-only controls, such that the media-only condition is mathematically centered at zero on the log2 ratio scale; deviations from zero therefore reflect stimulation-induced shifts in the relative balance of type 1 versus type 2 cytokines within individuals. The distributions of both raw and log2-transformed cytokine concentrations failed normality testing, so nonparametric tests were used for these analyses. For each cytokine, comparisons between the three treatment groups (SGE, LJM19, and LJL143) were performed with the Friedman test (due to having participant-matched samples) and Dunn’s test for multiple comparisons. To test for significant polarization towards type 1 or type 2 responses, we calculated the log2 of the ratio of each type 1 cytokine (IFN-γ or IL-12) to each type 2 cytokine (IL-4, IL-5, IL-10, or IL-13).^61^ Statistical analyses of cytokine ratios were performed on the log scale rather than the linear scale to stabilize variance and to mitigate extreme right-skews commonly seen in cytokine data distributions. To each ratio, we then applied the non-parametric Wilcoxon signed-rank test versus a value of zero as the null hypothesis (an equal balance of type 1 and type 2 cytokines) to determine whether the cytokine ratio was significantly skewed towards type 1 or type 2 immunity compared to cells treated with media alone.

For the *Leishmania* killing assay (Fig. 7), within each of the two exposure groups (*Lu. longipalpis* Unexposed or Exposed) pairwise differences were analyzed between PBMCs stimulated with media alone versus each of the other treatment conditions. For the percentage of infected macrophages, the dataset passed all normality tests with participant-matched samples, had significantly different variances between treatment groups, and had three missing samples (*n* = 1 from Unexposed/media and *n* = 2 from *Lu. longipalpis* Exposed/media due to random failure of these macrophages to adhere and grow in culture). Thus a mixed effects model was applied followed by Holm-Šídák’s test for multiple comparisons. To compare the percentage of infected macrophages between unexposed and exposed samples for LJM19 or LJL143, an unpaired t-test with Welch’s correction was used. The number of amastigotes per macrophage was found to have a lognormal distribution with no significant differences in variance between treatment groups. Thus, we used an unpaired lognormal t-test to compare the number of amastigotes per macrophage between unexposed and exposed samples for LJM19 or LJL143. Because the log2 type 1/type 2 cytokine ratios are non-normally distributed, we applied Spearman’s test to determine the correlation between the log2 ratio of type 1/type 2 cytokines and the percentage of infected macrophages. Correlation trend lines for non-parametric data were plotted using a smoothing spline with 3 knots.

## Acknowledgements

This work was supported by the NIH, Office of Clinical Research, Bench to Bedside Grant and was funded by the Division of Intramural Research, NIAID, NIH for S.K., J.G.V., and F.O. J.R.L. was supported by the NIAID Transition Program in Clinical Research. The funders had no role in study design, data collection, analysis, or interpretation, writing of the report, or the decision to submit the article for publication. The authors thank Dr. Mary Marovich for serving as the study monitor, Dr. Johannes Doehl for advice on statistical analyses, and the selfless volunteers who made this study possible.

## Author contributions

C.T., F.O., N.A., J.R.L., J.G.V., and S.K. conceptualized and designed the study. N.A., J.G.V., and S.K. wrote the human subjects protocol. R.R., K.H., G.W.T., and N.A. performed the clinical assessments and collected clinical samples. C.M. reared and provided the sand flies used for all human exposures and experiments. C.T., F.O., and S.K. performed the sand fly exposures on the study participants. M.A., C.T., R.D., R.G., W.C., and F.O. designed and conducted the experiments. M.A., C.T., F.F.A., N.A., J.R.L., J.G.V., and S.K. analyzed the data and performed statistical analyses. M.A., E.I., J.R.L., and S.K. wrote the manuscript. N.A., J.G.V., and S.K. supervised the study.

## Author disclaimers

This research was supported by the Intramural Research Program of the National Institutes of Health (NIH). The contributions of the NIH authors are considered Works of the United States Government. The findings and conclusions presented in this paper are those of the authors and do not necessarily reflect the views of the NIH or the U.S. Department of Health and Human Services.

The identification of specific products or scientific instrumentation is considered an integral part of the scientific endeavor and does not constitute endorsement or implied endorsement on the part of the authors, Department of Defense, or any component agency. The views expressed in this manuscript are those of the author and do not necessarily reflect the official policy of the Uniformed Services University of the Health Sciences, the Department of Defense or the U.S. Government.

Material has been reviewed by the Walter Reed Army Institute of Research. There is no objection to its presentation and/or publication. The opinions or assertions contained herein are the private views of the author, and are not to be construed as official, or as reflecting true views of the Department of the Army or the Department of Defense.

The investigators have adhered to the policies for protection of human subjects as prescribed in AR 70–25.

The findings of this study are an informal communication and represent the authors’ own best judgments. These comments do not bind or obligate the Food and Drug Administration.

## Declaration of generative AI and AI-assisted technologies in the writing process

During the preparation of this work, the authors used GPT-5.2 (OpenAI) in order to improve readability and language of the manuscript. After using this tool/service, the authors reviewed and edited the content as needed and take full responsibility for the content of the publication.

## Data availability

All data are available in the Supplementary Data File and are publicly available at https://doi.org/10.6084/m9.figshare.30397273.

