## Supplemental Figures and Table for "Human sand fly challenge elicits saliva-specific innate and type 1-polarized immunity that promotes *Leishmania* killing"

### SUPPLEMENTARY FIGURES AND TABLE

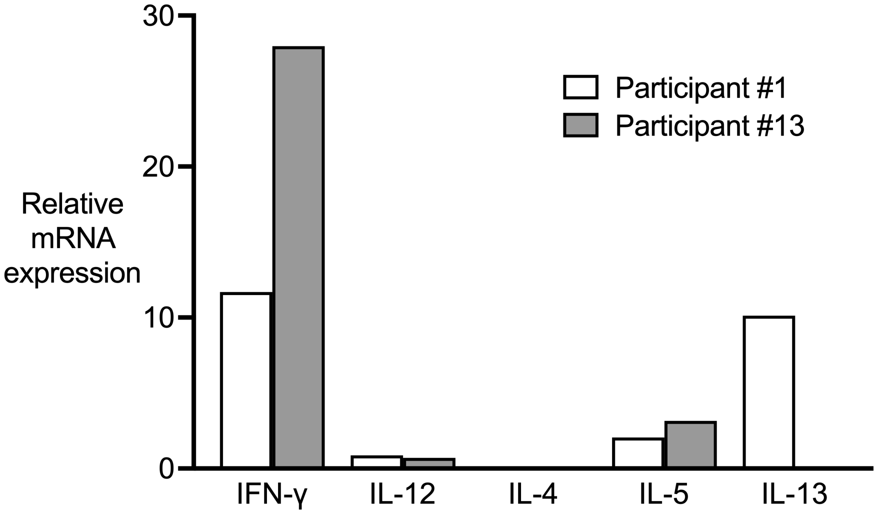


**Fig. S1.** Skin cytokine profiles of the delayed-type hypersensitivity response to *Lu. longipalpis* bites. Skin punch biopsies were collected as described in Fig. 3 for measurement of cytokine mRNA expression by quantitative RT-PCR. For each participant, gene expression at the bite site was normalized to expression in normal appearing skin from the contralateral arm.

**Supplemental Table 1.** PBMC batches by exposure number used for each experiment.

| Participant # | Figure 4 | Figures 5 and 6 | Figure 7 |
|---|---|---|---|
| **1** | 2 | 9 | 7, 8, 9 |
| **2** | 2 | 4 | - |
| **3** | 2 | 9 | - |
| **4** | 2 | 8 | - |
| **5** | 2 | 4 | - |
| **6** | 2 | 6 | - |
| **7** | 4 | 4 | - |
| **8** | 4 | 9 | - |
| **9** | 4 | 5 | - |
| **10** | 4 | 7 | - |
| **11** | 4 | 4 | - |
| **12** | 4 | 5 | 8 |
| **13** | 2 | 6 | 9 |
| **14** | 2 | 5 | 9 |
| **15** | 2 | 5 | - |

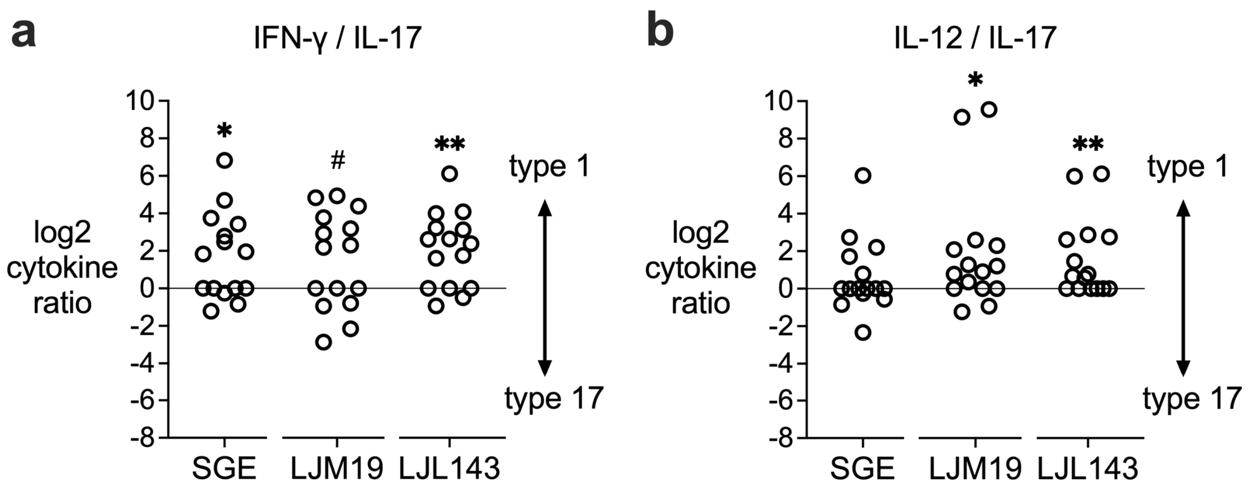


**Fig. S2.** LJM19 and LJL143 do not induce a type 17 cytokine response. PBMCs from sand fly-exposed study participants (*n* = 15) were stimulated with SGE, LJM19, or LJL143. Cytokine concentrations were measured by multiplex bead array and normalized to media-treated, participant-matched cells by background subtraction, as in Fig. 5. Ratios of the type 1 cytokines IFN-γ (**a**) or IL-12 (**b**) to IL-17 were calculated, where log ratios above 0 (solid line) indicate type 1 polarization while log ratios below 0 indicate type 17 polarization relative to media-treated cells, as analyzed by Wilcoxon signed-rank test using a value of zero as the null hypothesis (equal balance of type 1 and type 17 cytokines), # $p < 0.10$, * $p < 0.05$, ** $p < 0.01$.

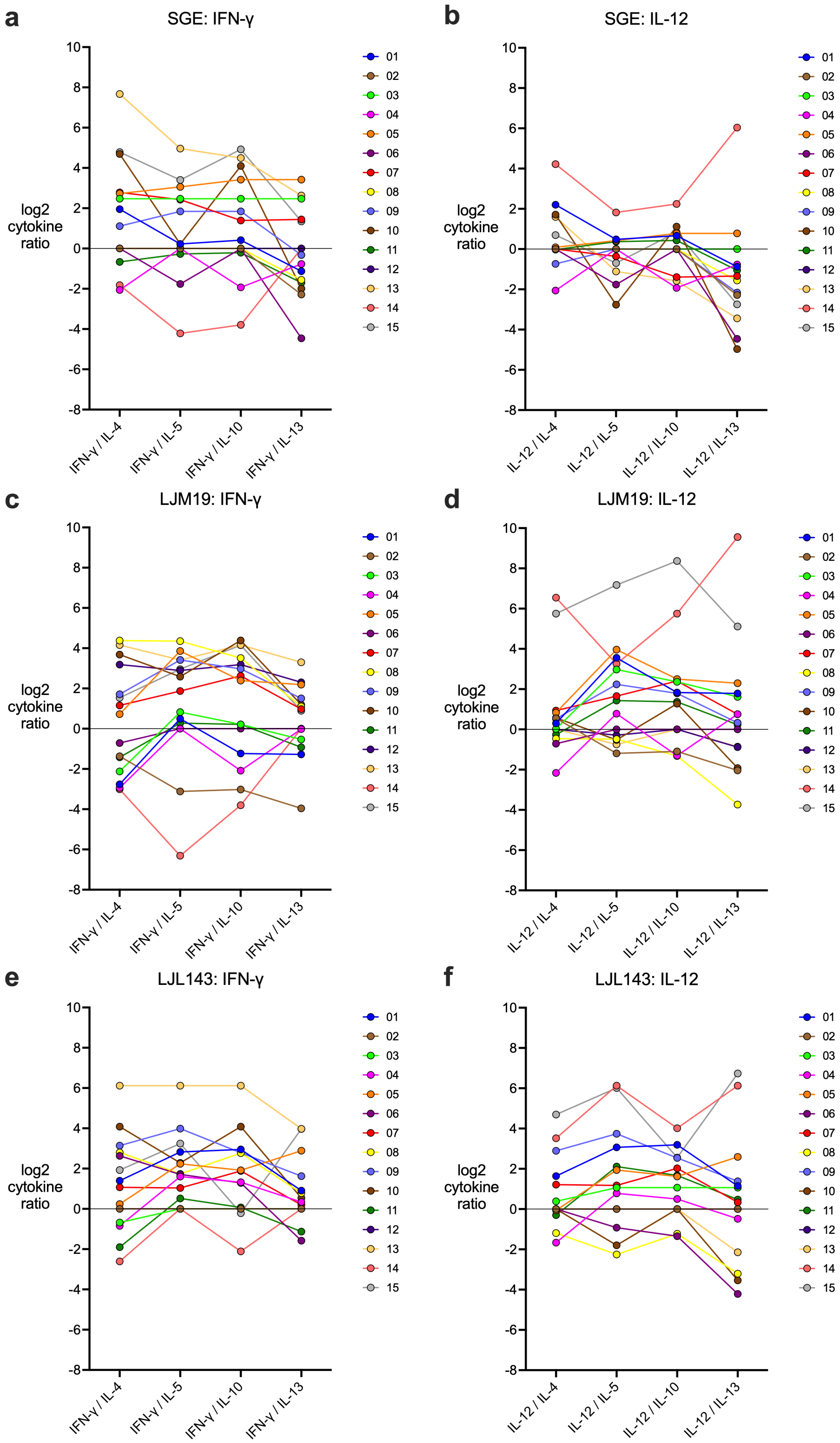


**Fig. S3.** Individual type 1/type 2 cytokine response profiles to *Lu. longipalpis* SGE and salivary proteins. Re-plot of data from Fig. 5 of cytokines produced by PBMCs from *Lu. longipalpis* exposed participants following stimulation with SGE (**a, b**), LJM19 (**c, d**), or LJL143 (**e, f**). Each color and line and their connecting points represent a single individual.

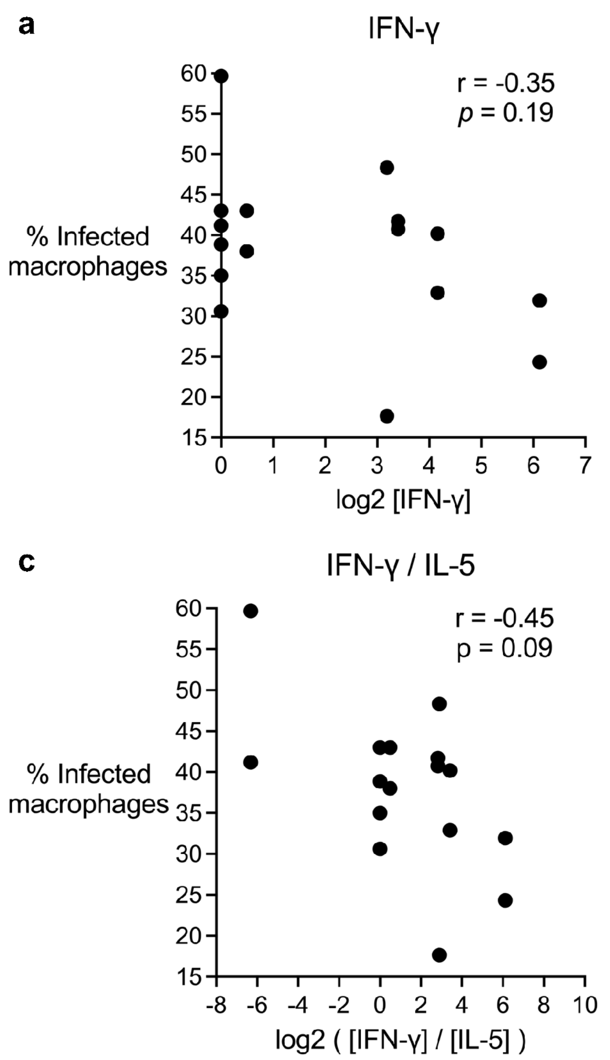


**Fig S4.** Correlation of $T_H$ cytokines with macrophage killing of *Leishmania* parasites. Spearman correlation between IFN-γ alone (**a**), IFN-γ/IL-13 ratio (**b**), or IFN-γ/IL-5 ratio (**c**) as calculated in Fig. 5 and the percentage of infected macrophages for LJM19- and LJL143-treated samples.
